# Gradients of local circuit architecture shape stability and information integration across the cortical hierarchy

**DOI:** 10.64898/2026.10.07.757185

**Authors:** Laura Green, Shinya Ito, Darrell Haufler, Marton Rozsa, Christine Wang, Kayvon Daie, Anton Arkhipov

## Abstract

The mammalian neocortex is a complex structure with many interacting regions and cell types. Multiple efforts have recently generated large datasets from multiple cortical regions to determine generalizing principles of cortical organization and activity. One such generalizing principle is the concept of a cortical hierarchy – low-dimensional macroscopic gradients of biological properties across cortical areas and a hierarchy of timescales for information processing. Such gradients in properties along a cortical hierarchy may explain how different cortical regions serve different computational roles (i.e. sensory processing versus working memory). We set out to investigate such principles using biologically realistic simulations. Starting from our previously published model of the mouse primary visual cortex (VISp) with updated connectivity reflecting recent large-scale connectomics and synaptic physiology datasets, we generated multiple versions of the cortical circuit model with increased connectivity as measured along the cortical hierarchy. We then measure the resulting neural activity in response to a simulated single-cell photostimulation paradigm and thalamocortical inputs. We found that overall circuit stability increased across the cortical hierarchy, due to both increases in connection density and increases in the relative proportion of SST cells. The results of this study link cortical cell types to population-level activity, generating insight into the function of local circuits across cortex.

## Introduction

The mammalian cortex is a complex system consisting of multiple areas that carry out a variety of incompletely understood functions, from sensory processing to higher-order cognitive operations, such as working memory, decision making, or language. These functions appear to be, at least in part, localized to certain areas in a hierarchical manner ^1–3^. A key question is how different aspects of cortical structure contribute to the function of a cortical region.

The computational function of each brain region is likely in part due to its connections to other brain regions. For example, visual cortex receives visual input from the retina via the thalamus, suggesting a role in processing visual information. Another example is motor cortex, which sends output directly to the corticospinal tract to control body movement ^4^. Other brain regions receive input from multiple sensory areas, suggesting a higher order, multimodal computational function such as working memory, decision making, or language. Supporting this idea, the hierarchical score of a cortical region is typically calculated as the ratio of feedforward to feedback input to a region, such that regions that receive more feedforward input from a large number of areas are considered higher in the cortical hierarchy and are known to be more involved in higher order cognitive function ^1,5–7^. Another method to determine the function of a region is to look at innervation from subcortical nuclei known to be important for neuromodulator function^8–10^.

However, in addition to global brain properties that vary along the cortical hierarchy, there are also variations in local circuit properties, such as neuron density^11,12^, synaptic density^11,13^, synaptic protein gradients^14,15^, the extent of dendritic branching^16–18^, pyramidal cell genetic variability^19,20^, neuromodulator receptor density^14,21,22^, cytoarchitecture^23^, and the ratio of parvalbumin (PV)-positive versus somatostatin (SST)-positive inhibitory neurons^24,25^. There are two best-studied properties that change across the cortical hierarchy. The first is a change in the relative number of inhibitory neuron subtypes, such that there is a higher density of PV neurons in primary areas than higher order areas, while the opposite is true for SST neurons ^24,25^. The second property is the number of synapses and extent of dendritic branching across the cortical hierarchy, such that higher order areas likely have higher connection density than primary areas. This is especially true in human and non-human primates, but is demonstrated more subtly in rodents as well ^13,26–32^. A major question therefore is how these local gradients support the functional hierarchical organization.

Recent technological advances and organized efforts have enabled the collection of increasingly large and detailed datasets in neuroscience^24,33–38^, offering excellent opportunities to study organizational principles of the brain. Computational simulations provide a powerful framework for making use of these data. They allow detailed measurements of cellular and circuit properties to be integrated into models that can be used to investigate contributions of these properties to circuit activity. Here, we applied this approach to study the relevance of structural gradients across hierarchical areas to cortical function. We constructed biologically-realistic models representing five cortical areas across the cortical hierarchy, integrating connectivity and cell types information to investigate how these features shape neural activity^39–42^. While simulations at the whole cortex level have been demonstrated^43^, they require immense computational resources, and comprehensive datasets needed to constrain models at the scale of the entire cortical hierarchy are not yet available. Therefore, this work focused on studying local circuit properties while modeling multiple cortical areas across the hierarchy.

The circuit models we constructed varied either in connection density, inhibitory neuron type density, or both, so that we could assess the effects of each gradient on model activity independently and in conjunction. This was feasible due to recently published datasets on the synaptic properties and connectome of the mouse primary visual cortex (VISp) ^34,38,44^, as well as recently published large-scale models ^39,45^. The circuit models presented here contain all cortical layers, diverse excitatory and inhibitory cell types, experimentally measured neuronal densities, neuron and synapse type models capturing experimentally observed electrophysiology, and connectivity systematically derived from synaptic physiology and connectomics data. These features facilitate exploration of the relations between circuit structure and function under biologically realistic We measured the differences in firing rate, excitatory-inhibitory balance, time constants, and response to stimulation across the models. In this way, we were able to study how gradients in local properties contribute to differences in function across the cortical hierarchy.

We found that firing rates, stimulus selectivity, and excitatory-inhibitory balance varied systematically with local circuit properties. Increasing connection density and SST neuron density enhanced excitatory-inhibitory balance and stability of cortical circuits in response to perturbations, with SST neuron density in particular being important for maintaining stability in higher-order models. We also found that increasing SST density was associated with an increase in convergent network motifs, suggesting a potential role for SST neurons in integrating inputs in higher-order cortical areas. In contrast, local changes in connection density and inhibitory neuron composition did not account for the increase in neural timescales observed across the cortical hierarchy, suggesting that this gradient may arise from factors beyond the local circuit. Finally, single-cell stimulation revealed strong recruitment of SST neurons, arising from both their intrinsic excitability and relatively weak inhibitory input, and provided a direct link between circuit structure and cell-type-specific responses. Overall, we demonstrate the efficacy of biologically-realistic large-scale modeling in connecting detailed structural data to the function of a circuit.

## Results

### SST input to excitatory cells increases across the cortical hierarchy

We created 5 biologically-realistic models of circuits along the cortical hierarchy, to study the relation between relative inhibitory neuron density and connection density on circuit function. These models were based on a previously published biorealistic model of the primary visual cortex, fit to publicly available large-scale connectomic and multi-patch physiology datasets ^39,45^. They consisted of point neurons (generalized leaky integrate-and-fire) representing 19 major cell types (PV, SST, VIP, as well as excitatory cells corresponding to each cortical layer), fit to reproduce spiking activity from electrophysiological data ^46^. Neurons were arranged into 5 layers (L1, L2/3, L4, L5, L6) with distance-dependent connectivity derived from electron microscopy and synaptic physiology datasets^34,38^. Synaptic weights and delays were derived from a large-scale multi-patch physiology dataset^34^.

We chose to model two primary regions (primary visual cortex and primary motor cortex) and three higher order regions (posterior parietal cortex, anterior cingulate cortex, and orbitofrontal cortex) that represented a gradient in inhibitory neuron density and cell density (Fig. 1a-d. The model representing a sensory area at the bottom of the cortical hierarchy had 65,874 neurons and 23,773,150 connections, while the model representing an area at the top of the cortical hierarchy had 21,968 neurons and 22,692,023 connections. The goal when generating these models was to decrease the neuron count while maintaining the same number of connections, such that connection density increased along the cortical hierarchy.

**Figure 1.**
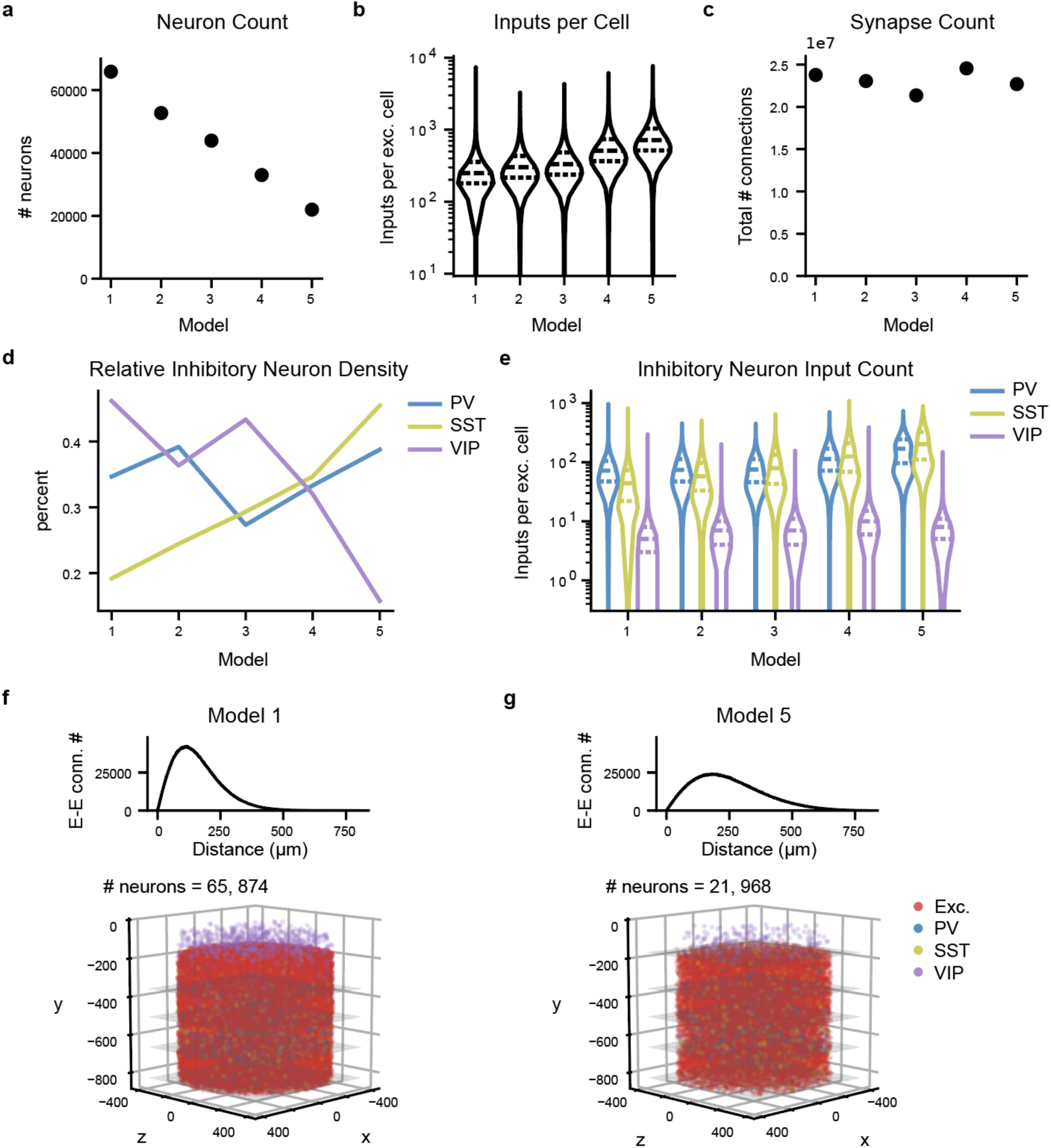
Circuit properties across the cortical hierarchy. a: The number of neurons represented in each model along the cortical hierarchy, informed by the density of neurons under the surface area of the cortex. b: The total number of synapses in each model. c: The total number of inputs per excitatory cell in each model. d: Percentage of each of the three major inhibitory neuron subclass in each model (data adopted from ^24,25^). e: The number of inputs per excitatory cell from each inhibitory neuron subclass in each model. f: Representation of the network of model 1. Top: Number of connections between excitatory cells at a range of distances between neurons. Bottom: Representation of the neurons within the model, colored by cell type category. g: Same as (f), but for model 5.

The models received input from a thalamus-like input model, consisting of 17,400 virtual spike-generator units modelled as filters and fit to *in vivo* electrophysiology data from cells with diverse response types in the thalamus (see Methods and ^39^ for more details). The thalamic input to each cortical model was the same, so that any variations in the output of the cortical models could be attributed to the variations in connectivity and inhibitory neuron density.

The neuronal density decreased in models along the cortical hierarchy (Fig. 1a) while the number of inputs to each cell increased (Fig. 1b). The total number of synapses remained constant, with some variability due to the stochastic nature of connection generation in our model building procedure (Fig. 1c; see methods).

We varied the density of PV, SST, and VIP neurons according to data from ^24,25^ (Fig. 1d). As a result, our first observation is that excitatory cells in the models at the bottom of the cortical hierarchy receive more PV inputs than SST inputs, while excitatory cells at the top of the cortical hierarchy receive more SST inputs than PV inputs (Fig. 1e). The average number of VIP inputs remains relatively constant, because even though the proportion of VIP neurons decreases, the overall number of synapses in the model increases.

Besides the 5 models that vary in both connection density and inhibitory neuron subtype density, we created an additional 4 models that vary in connection density alone and 4 models that vary in inhibitory density alone. The goal of these additional models was to parse whether changes in output of each network along the cortical hierarchy are due to connection density, inhibitory neuron subtype density, or both. The result was 3 groups of 5 models, with the same lowest order cortical area model in each group, thus resulting in 13 total unique models.

### Population-level activity is qualitatively similar across models

Our first test was to provide a standard stimulus to each of our models across the cortical hierarchy and validate the neural activity observed in simulations of these models against experimental data. Qualitatively, the output of each model was similar, with overall asynchronous spiking activity and increased activity in response to stimulus onset (Fig. 2a, b), similar to our previously published simulations of mouse primary visual cortex (VISp) ^39,45^. The proportion of active cells was lower in the higher order than in the lower order models, likely due to the decrease in weights proportional to the increase in connection density in the higher order model (Fig. 2b; see Methods). Similarly, both spontaneous and evoked firing rates decreased across the cortical hierarchy in layers 4-6 (Fig. 2c, d). In the models where we did not vary the connection density, we do not observe a decrease in firing rates (Fig. S1).

**Figure 2.**
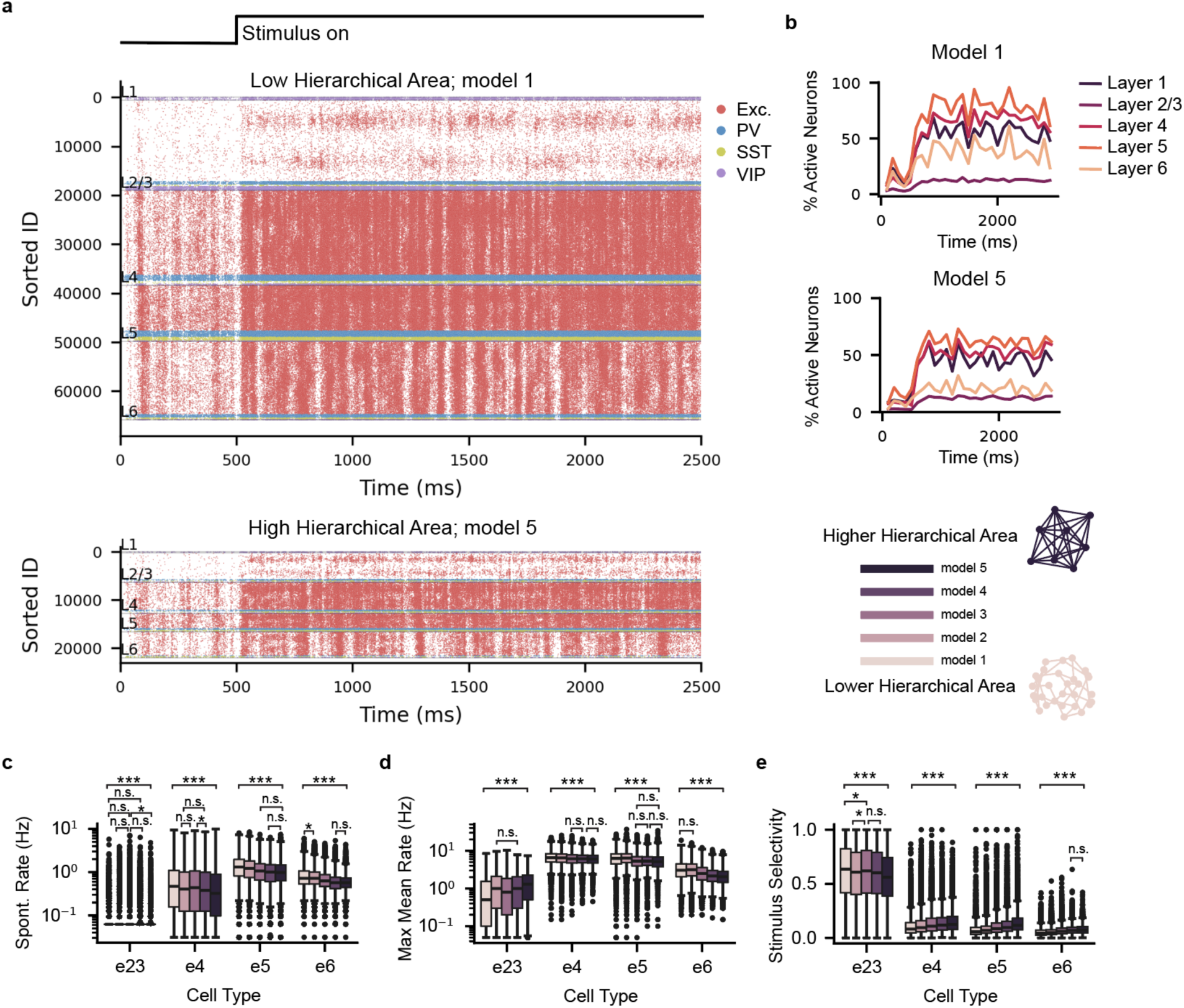
Activity of cortical circuit models across the cortical hierarchy. a: Activity of cortical circuit models in response to increased input, in a model lowest in the hierarchy (top) and a model highest along the cortical hierarchy (bottom). Response of all neurons in the models to a realistic excitatory stimulus presented at t=500 ms. b: Proportion of active neurons in each layer, measured as the percentage of neurons that spike at least once in each 100 ms bin during the trial. Top: the lowest hierarchical model, as shown in (a, top); Bottom: the highest hierarchical model, as shown in (a, bottom). c: Firing rates during spontaneous activity of excitatory cells in layers 2/3, 4, 5, and 6 (“e2/3”, “e4”, “e5”, and “e6”, respectively) along the cortical hierarchy. Darker colors correspond to higher order cortical areas. d: Firing rates during stimulus presentation of excitatory cells in layers 2/3, 4, 5, and 6 along the cortical hierarchy. e: Stimulus selectivity of excitatory neurons in layers 2-6 along the cortical hierarchy.

We found that stimulus selectivity was higher across the cortical hierarchy, even when controlling for the decreased firing rates (one-way ANCOVA, F(4, 39206)=776.1, p<0.001). We also found a decrease in normalized power in lower frequency bands (5-15 Hz) and increase in high frequency bands (60-100 Hz) across the cortical hierarchy in deeper layers, but only in the models with both connectivity and inhibitory neuron density changes (Fig. S2), generally matching trends in large-scale physiology datasets^47^.

### Excitatory-inhibitory balance increases across the cortical hierarchy

To study why firing rates decreased across the cortical hierarchy in deeper layers but not superficial layers, we next examined how increasing the connection density and inhibitory type density differentially recruit excitatory versus inhibitory activity in each model. We computed the total input to excitatory cells from either other excitatory cells, inhibitory cells, or external input during the stimulus (Figure 3a) and measured excitatory/inhibitory balance by the ratio of total input to excitatory input (from both recurrent and external sources), denoted by the metric ß ^48^.

**Figure 3.**
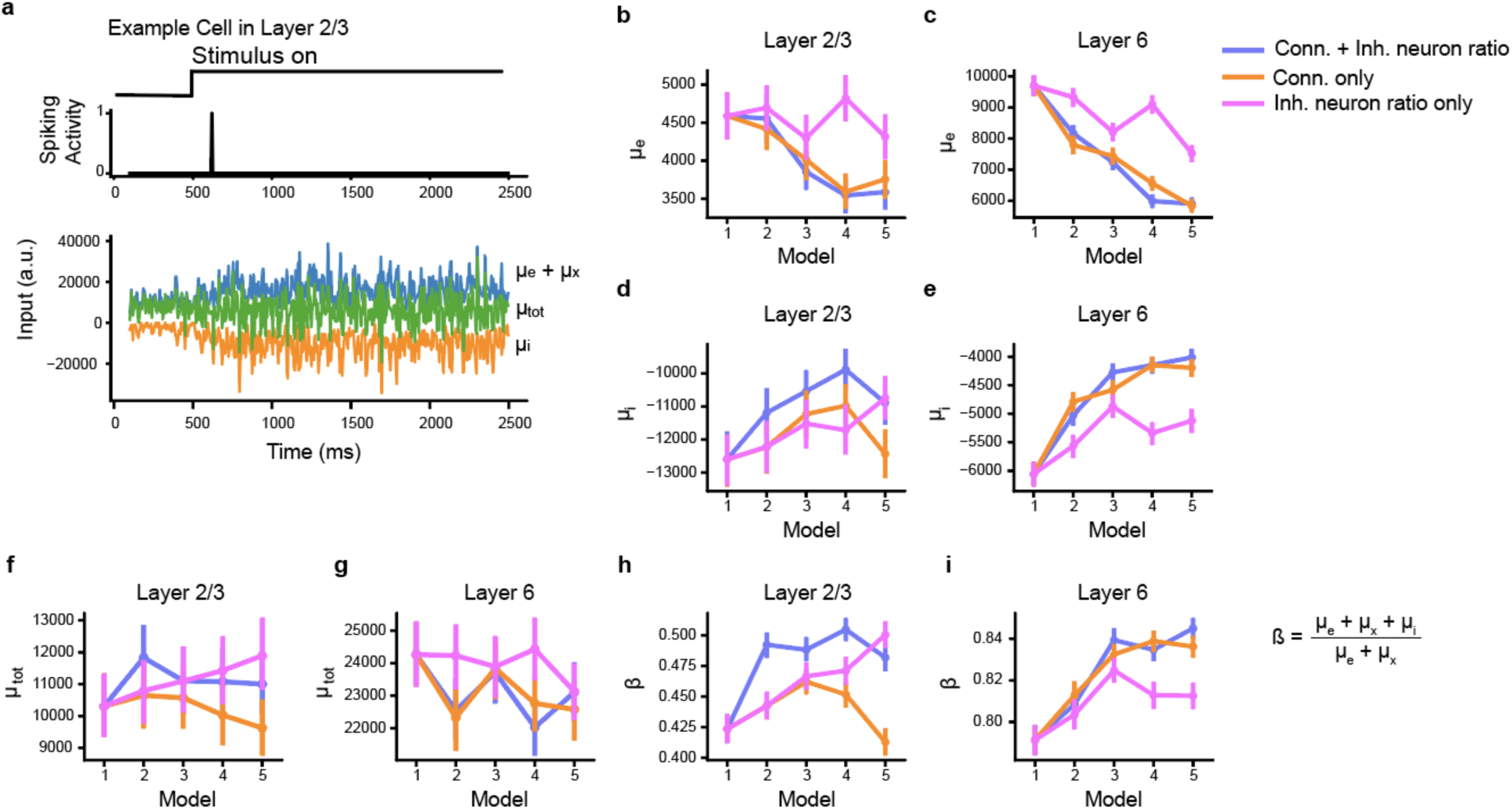
E/I balance increases across the cortical hierarchy, due to weakening recruitment of local input. a: Spiking activity of a L2/3 excitatory cell during a single trial (top) and input to the same cell (bottom). Blue: excitatory input (local excitatory input (µ_e_) and external input (µ_x_)); orange: inhibitory input (µ_i_); green: total input (µ_tot_). b, c: Recurrent excitatory input to excitatory neurons in L2/3 (b) and L6 (c). d, e: Inhibitory input to excitatory neurons in L2/3 (d) and L6 (e). f, g: Total input to excitatory neurons in L2/3 (f) and L6 (g). h, i: Balance metric, ß, for L2/3 (h) and L6 (i) excitatory neurons across all 13 model types.

Excitatory and inhibitory input to excitatory cells decreased across the cortical hierarchy in all layers and model types (Fig. 3f-i, also see S3 for layers 4 and 5), suggesting that both changes in connection density and inhibitory neuron type density lead to rebalancing in cortical circuits. However, the relative decrease in excitatory input and inhibitory input varied across the hierarchy. The decrease in excitatory input was greater than the decrease in inhibitory input in models with connection density changes only, such that decreasing connection density led to decreased total input to excitatory cells in both layers 2/3 and layer 6 (orange line in Fig 3d, e, also see Fig. S3 for layers 4-5). This was likely due to the decreased weights in the higher order model. Increasing the SST neuron density led to increased total input to layer 2/3 neurons, but not to layer 6 neurons. Changing both connection density and inhibitory neuron type density led to no change in total input in layer 2/3, and a decrease in total input in layer 6, which explains why firing rates decreased in higher order models in deeper layers but not superficial layers (Figs. 2d, 3d, e).

We next calculated ß as a measure of how close a neuron is to a fluctuation driven regime (when excitatory input is of the same magnitude as the total input and evenly balanced by inhibitory input) or an input driven region (when large changes in excitatory input are required to drive activity in a neuron, because mean excitatory input is much lower than the mean inhibitory input). We found that ß was high (>0.1) across all models and increased across the cortical hierarchy, indicating that the primary sensory model was in a fluctuation-driven regime, and that increasing connection density and inhibitory neuron subtype density led to an increasing shift towards the fluctuation-driven regime. The value of ß varied between layers, such that layer 6 had the highest ß value, followed by layer 4, layer 2/3, then layer 5.

### Connection density and inhibitory neuron density do not contribute to longer timescales across the cortical hierarchy

Experimental studies suggest that time constants of neural activity tend to increase along the cortical hierarchy. This is thought to be associated with the requirements for fast sensory processing in primary areas versus the accumulation of contextual information in higher order areas ^49,50^. This observation has been replicated across species—from humans ^50,51^, to non-human primates ^52,53^, to rodents ^33,54^, using a range of recording techniques. Previous work poses a question whether the gradient in timescales may be due to increased recurrence or variation in inhibitory gene expression ^51,54,55^. Here we examined whether increased recurrence and inhibitory neuron subclass ratio in local circuits contribute to an increase in timescales across the cortical hierarchy.

We measured the intrinsic timescale of a circuit by taking the autocorrelation of the activity of individual excitatory neurons and measuring the time to half of maximum amplitude (Fig. 4a). The time constants ranged from 50 to 90 ms across layers and models, consistent with intrinsic time constants observed *in vivo* ^33,54,56^. However, we found that increasing connection density led to a slightly shorter, not longer, time constant across layers (Fig. 4b-c, also see Fig. S4). We also found that the time constant was longer in layer 6 than in layers 2/3 by approximately 60% or 30 ms, and since layer 6 has denser recurrent connectivity than layer 2/3, recurrence may still contribute to longer time constants in the cortex. Changing the relative inhibitory neuron density had no effect on time constants in layer 2/3 but did lead to a small decrease in the time constant in layer 6.

**Figure 4.**
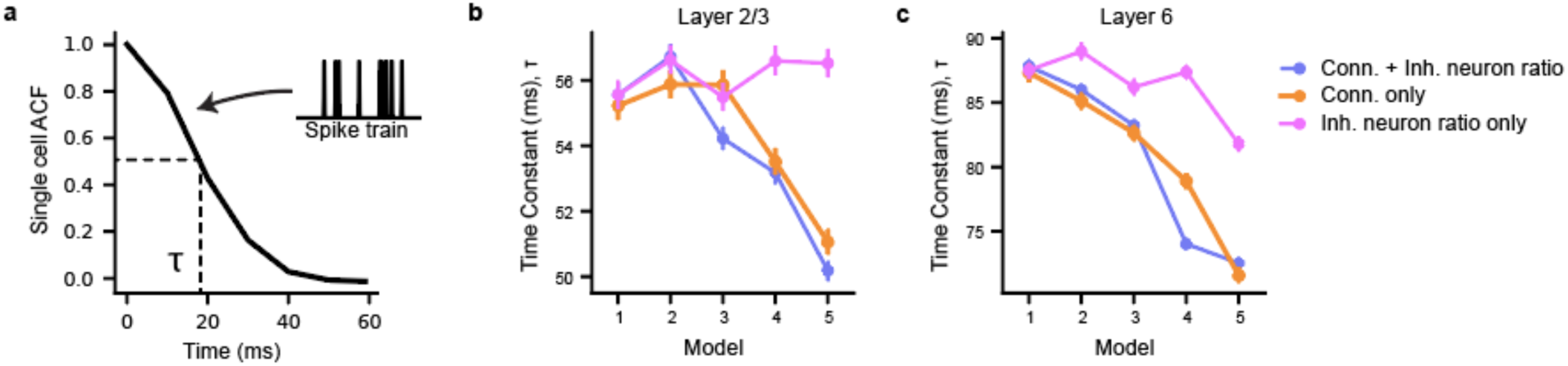
Population dynamics across the cortical hierarchy a: Autocorrelation time constants for each neuron are calculated as the time to half of the peak of the autocorrelation function of a cell’s spiking activity. b: Time constants of single neuron activity across the cortical hierarchy in layer 2/3. c: Same as (b), but for layer 6.

The decrease in time constants across cortical hierarchy models suggests that local circuit properties are not responsible for the increase in time constants observed *in vivo*. Indeed, previous work showed a lack of correlation between timescales of individual neurons within a region during different behavior epochs, indicating that the timescale gradient is likely due to a factor that varies independently within populations of neurons in each brain area ^54^. Other potential mechanisms may include increased feedforward connectivity and variations in neuromodulator expression.

### Small world motifs, especially convergent motifs, increase across the cortical hierarchy

We next examined the effects of changing the network structure on the graph properties of the network. It has been shown that neurons with similar tuning properties are more likely to connect to each other and share stronger connection strengths^39,57–59^, which we replicated in our models as well (see Methods). We expected that increasing connectivity, combined with the like-to-like rules in the network, would lead to an increase in the clustering coefficient and small world properties of the network.

Increasing the relative density of inhibitory neuron types could also potentially affect the graph structure of the network, since the connection profile is different between inhibitory neuron types ^34,38,44^. SST neurons project fewer outputs than PV neurons but receive a greater proportion of inputs from excitatory neurons versus other types of neurons (Fig. S8) ^44^.

Increased clustering in small world networks is associated with signal-propagation speed, computational power, and synchronizability, but also reduced dynamic stability ^60^. However, the prevalence of highly interconnected subgraphs within a network can improve local dynamic stability, with consequences for the entire network ^61–63^. Indeed, sparse reconstructions of networks across the mouse cortex demonstrated an increase in motif count across the cortical hierarchy and suggested a role in enhancing dynamical stability ^64^. We thus measured the change in both clustering coefficient as well as the prevalence of highly connected motifs within each of our network models to study the change in graph properties of cortical regions across the cortical hierarchy.

We found that the clustering coefficient increases across the cortical hierarchy (Fig. 5a, b, and see Fig. S5 for layers 4-5) with increasing connection density, but not with increasing SST density, in both layers 2/3 and 6 (Fig. 5c). In layer 6, increasing inhibitory neuron type density led to a slight decrease in clustering coefficient. Clustering coefficients ranged from approximately 0.05 to 0.35, which is comparable to the clustering coefficients measured in *Drosophila Melanogaster*, 0.0463, and in layer 2/3 of mouse VISp, 0.159 (Lin et al., 2024).

**Figure 5.**
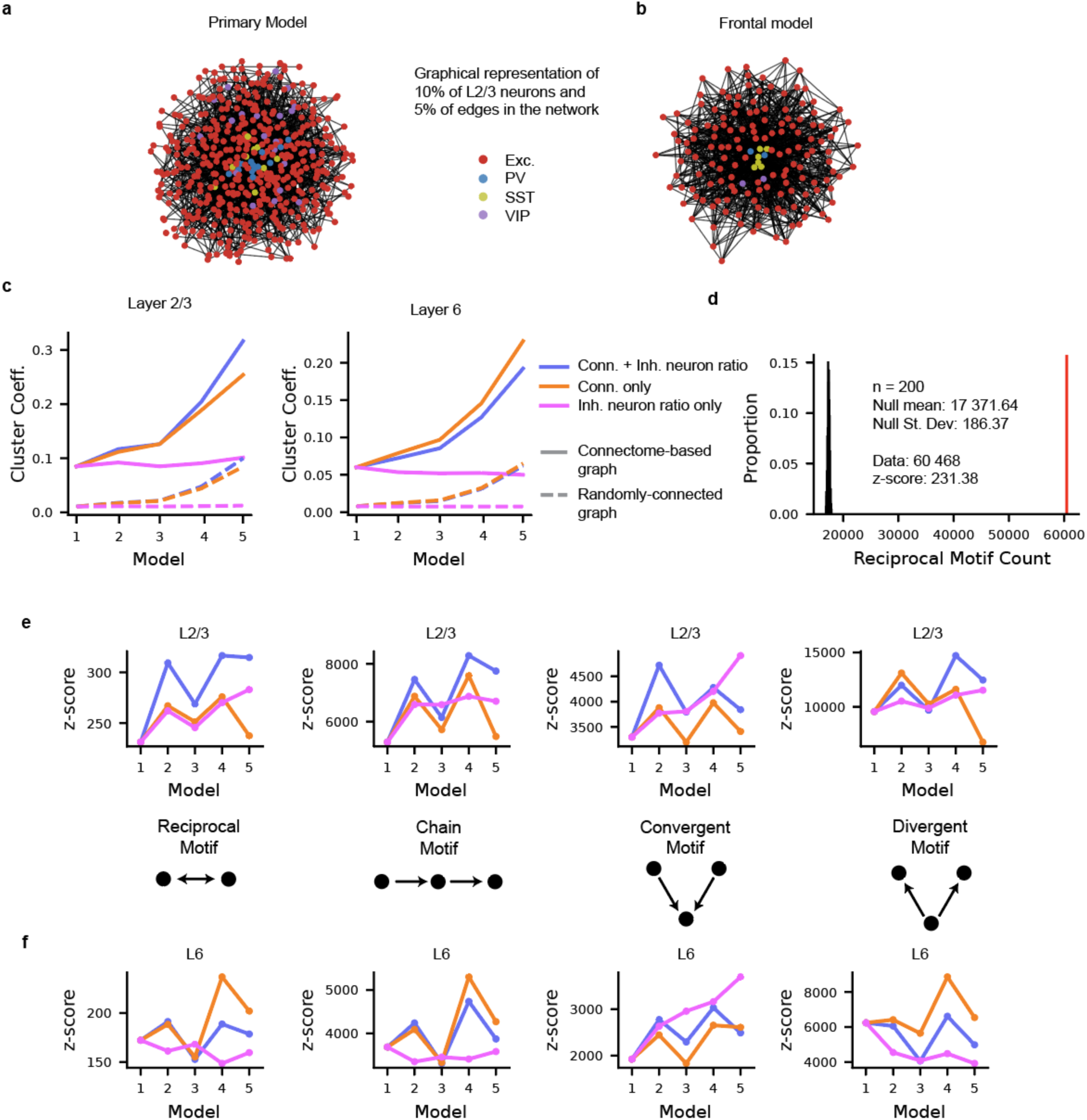
Small world network characteristics increase across the cortical hierarchy. a: graphical representation of model 1, representing a lower hierarchical region. b: Same as (a), but for model 5, representing a high hierarchical region. c: Cluster coefficient of the network across the cortical hierarchy in layer 2/3 (left) and layer 6 (right). Solid lines: Measurements from the models, based on the mouse connectome; dashed lines: measurements from an Erdős-Rényi null graph with the same number of nodes and edges. d: Number of reciprocal motifs in layer 2/3 of the lower hierarchical model (red line), compared to the number of reciprocal motifs in 200 Erdős-Rényi null graphs with the same number of nodes and edges (histogram). The overrepresentation of reciprocal motifs is measured by the z-score of the motif count relative to the mean and standard deviation of the null distribution. e: Overrepresentation of all motifs with 2 edges in L2/3 of all cortical models. f: Same as (e), but for L6.

We also examined the prevalence of all motifs with 2 edges – recurrent connections, chains, divergent connections, and convergent connections. In general, all motifs were highly over-represented in the cortical circuit models relative to random Erdős-Rényi networks, measured by the z-score of the number of motifs in the cortical circuit model relative to the mean and standard deviation of the number of motifs in 200 instantiations of a random network with the same number of nodes and edges (Fig. 5d-f). There was a trend of increasing prevalence of reciprocal and chain motifs across the cortical hierarchy, associated with an increase in the prevalence of reciprocal and chain motifs in the models with increasing inhibitory neuron subtype density in layers 2/3 (Fig. 5e). The trend was unclear in layer 6 (Fig. 5f).

There was an increase in convergent motifs across the cortical hierarchy with increasing inhibitory neuron subtype density in both layers 2/3 and layer 6, and either a decrease or no increase in divergent motifs across the cortical hierarchy (Fig. 5e-f). The increase in convergent motifs aligned with our prediction that increased SST neuron density increases the prevalence of convergent motifs and may play a role in routing or reducing the dimensionality of a larger number of feedforward inputs to higher order cortical areas ^24,65^.

### Inhibitory stabilization increases across the cortical hierarchy

Recent work has explored population dynamics across cortical layers and regions through inhibitory stabilization (Fig. 6a,b), which is indicative of strongly recurrent networks ^66–71^. Activation of inhibitory populations in strongly-recurrent networks leads to a paradoxical effect, where the firing rate of inhibitory populations decreases in response to their activation. This decrease in inhibitory activity is due to a decrease in local excitatory input following the initial brief stimulation-induced increase of inhibitory activity. At stronger levels of activation, inhibitory firing rates begin to increase (Fig. 6b). Conversely, activation of inhibitory neurons in weakly-connected networks does not lead to a paradoxical effect. The firing rate of inhibitory populations increases, while the firing rate of excitatory populations decreases (Fig. 6a).

**Figure 6.**
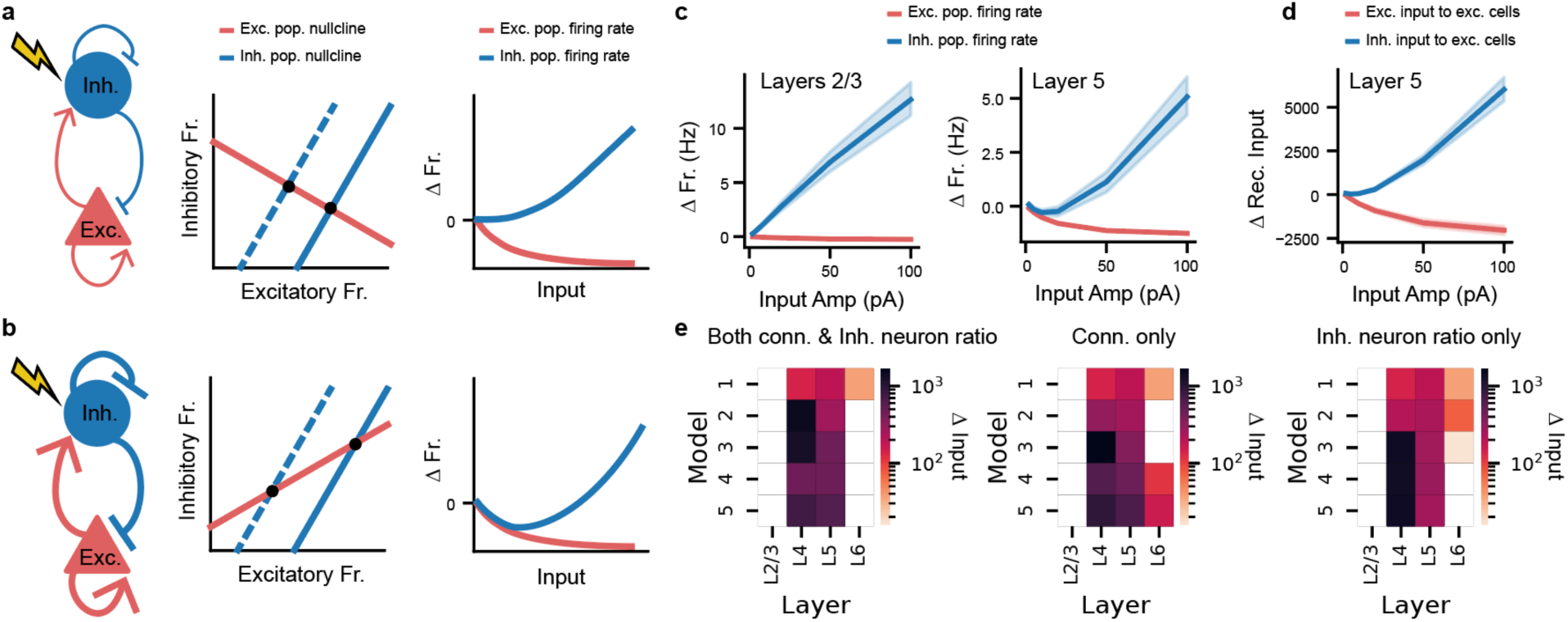
Inhibitory stabilization across the cortical hierarchy. a: Description of a weakly-coupled inhibitory-excitatory network. Left: schematic of weakly coupled inhibitory and excitatory populations. Middle: nullclines of the excitatory (red) and inhibitory (blue) populations. Black circles are the fixed points. Right: resulting firing rates of excitatory (red) and inhibitory (blue) populations with increasing input to inhibitory cells. b: Description of the paradoxical effect in a recurrent network with inhibitory stabilization. Left: Schematic of strongly coupled inhibitory and excitatory populations with strong recurrent weights. Middle and right panels are the same as (a), now showing decreasing inhibitory firing rates in response to a range of increased input to inhibitory cells. c: Firing rates of excitatory and inhibitory populations in a low hierarchical model (model 1) in response to increasing current input to all inhibitory neurons in the model. Left: response in layer 2/3. Right: response in layer 5. d: Change in excitatory input to excitatory and inhibitory neurons in layer 5 of the low hierarchical model. e: The maximum mean change in inhibitory input to excitatory cells while providing depolarizing current input to all inhibitory neurons in each layer and each model across the cortical hierarchy. Left: models with changes in both connection density and inhibitory neuron subtype ratio; middle: models with changes to connection density only; right: models with changes in inhibitory neuron subtype ratio only.

Since we modelled increasing recurrence across the cortical hierarchy, it follows that inhibitory stabilization may increase across the cortical hierarchy as well. To test this, we provided depolarizing current to all the inhibitory cells in each model at increasing intensity levels. We found that paradoxical responses occurred in layer 5 in lower-hierarchy model in response to inhibitory activation, but not in layer 2/3: Namely, the activity of inhibitory neurons decreased in response to increasing depolarizing input of amplitudes 0-25 pA in layer 5, but not in layer 2/3 (Fig. 6c). The paradoxical response in layer 5 was associated with increased inhibitory input to excitatory cells, confirming that the paradoxical response is due to inhibitory stabilization (Fig. 6d) ^72^. Inhibitory input to excitatory cells in layer 5 during inhibitory stimulation increased with both connectivity and relative SST density, though connectivity had the strongest effect (Fig. 6e).

### Single cell stimulation recruits superficial SST neurons

We next studied the relation between circuit structure and the function of specific cell types within each model. A recent method used to connect circuit structure to function is single-cell stimulation, in which experimentalists activate a single neuron at a time *in vivo* and measure the resulting effects on the surrounding neurons ^58,71,73–75^. We replicated the single-cell stimulation paradigm in our models and directly compared the results to experimental data, finding that our results matched the trends in the data overall.

Experimentally, we used a laser to excite single excitatory neurons optogenetically in the mouse primary motor cortex (MOp). The stimulus duration was 100 ms with 1 second intervals between stimulation periods. All other neurons were imaged simultaneously using 2-photon calcium imaging. In the models, we replicated this experiment by stimulating 55 excitatory neurons in layer 2/3 with a current injection, one neuron at a time (Fig. 7a). Each neuron was stimulated for 100 ms, a total of 25 times, with 1 second intervals between stimulation periods. The current injection was calibrated to lead to at least 10 extra spikes during the stimulation period.

**Figure 7.**
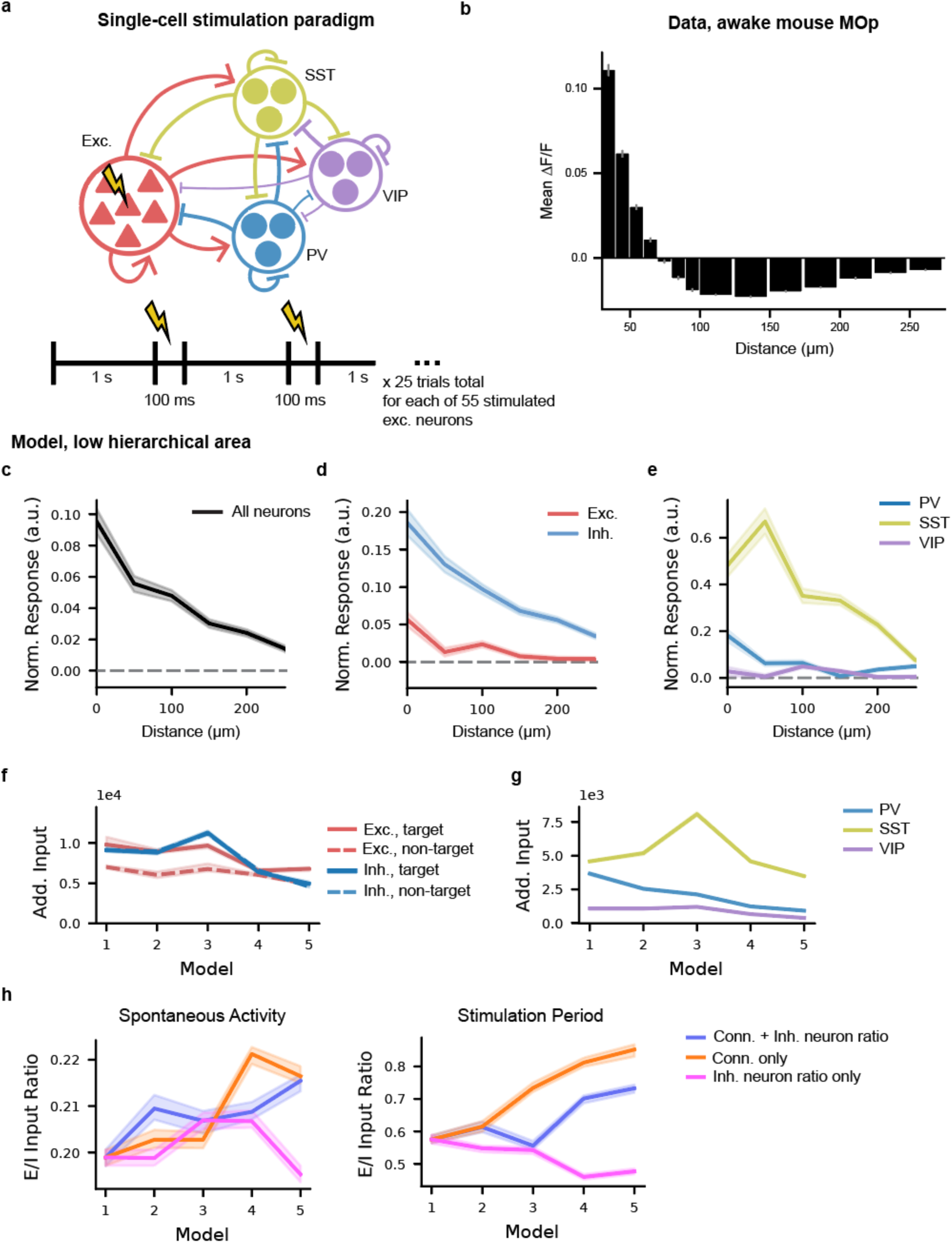
Single-cell stimulation of excitatory neurons *in vivo* and *in silico*. a: Schematic of the single cell stimulation experimental paradigm. b: Response of all measured cells within 250 µm of the stimulated neuron *in vivo* in mouse primary motor cortex (MOp). c: Response of layer 2/3 neurons in models across the cortical hierarchy during the single-cell stimulation paradigm. Activity is z-scored to the mean and standard deviation of activity during control trials without stimulation. Right: Excitatory cells. Left: Inhibitory cells. d, e: Same as (c), but separated by excitatory and inhibitory cells (d), and by inhibitory cell subclass (e). f: Change in excitatory (red) and inhibitory (blue) input to excitatory cells during the stimulation paradigm. Solid lines: input to cells receiving direct connections from the stimulated neuron; Dashed lines: input to cells not receiving direct connections from the stimulated neuron. g: Change in inhibitory input to excitatory cells by inhibitory cell type: PV (blue), SST (yellow), and VIP (purple). h: Ratio of excitatory to inhibitory input to excitatory cells across the cortical hierarchy for each model type during spontaneous activity (left) and the stimulation paradigm (right).

To measure the effect of single-cell stimulation, we analyzed how neurons throughout the cortical network responded to the stimulation depending on their distance to the stimulation site. *In vivo*, cells close to the stimulated neuron responded strongly, while cells further away responded weakly (Fig. 7b). The experimental paradigm did not allow for separation of responses by cell type. In our model, the average population response across layer 2/3 was also strong in the center and weaker further away, matching the experimental data (Fig. 7c). We could then further analyze the responses in the model by cell type to provide insight into the experimental data and circuit function within the model.

When looking at the responses of excitatory cells alone, we found that excitatory cells in layer 2/3 demonstrated a decreasing response to single-cell stimulation with distance to the stimulated neuron (Fig. 7d). In higher order models, excitatory neurons were only weakly excited, or even inhibited, by the single-cell stimulation paradigm (Fig. S6). On average, inhibitory cells were more strongly activated than excitatory cells and more responsive at increasing distances from stimulation (Fig. 7d). Across the cortical hierarchy, average responses of inhibitory neurons remained high (Fig. S6).

When we grouped inhibitory responses by inhibitory neuron subclass, we found clear differences in the responses of each subclass. SST neurons responded strongly while PV cells responded weakly near the stimulated excitatory neuron and not at all at further distances (Fig. 7e), similar to experimental observations ^73,76^. VIP cells were only weakly responsive to this stimulation paradigm (Fig. 7e). SST and VIP activation remained consistent across the cortical hierarchy, while the activation of excitatory neurons and PV neurons decreased across the cortical hierarchy (Fig. S6).

We next examined the change in input to excitatory cells in layer 2/3 due to stimulation, measured as the incoming spikes to each cell multiplied by the weight of each connection. All cells underwent an increase in both excitatory and inhibitory input during the stimulation period, on average (Fig. 7f). Cells that had direct connections (“target”) from the stimulated neuron received more excitatory input than cells that had no direct connections (“non-targets”), but both target and non-target cells received the same amount of inhibitory input on average. The overall effect was broad silencing or sparse responses in most excitatory cells, while a small number of excitatory neurons near the stimulated neuron responded more strongly (Fig. 7d, Fig. S6). SST neurons were responsible for most of the increase in inhibitory input during the stimulation paradigm, and SST input remained consistent in models across the cortical hierarchy, despite decreasing inhibitory input from PV and VIP neurons (Fig. 7g). In models without an increase in SST neuron density, SST input and overall inhibitory input to excitatory neurons decreased in higher order cortical models, leading to a larger increase in excitatory input relative to inhibitory input (Fig. S7).

We hypothesized that the high responsiveness of the SST neurons over the PV neurons could be due to either increased excitatory input, decreased inhibitory input, or intrinsic excitability of the SST neurons. We tested these hypotheses by measuring the excitatory and inhibitory input to SST and PV neurons in layer 2/3, and by swapping our SST neurons with PV neurons in the model. We first found that SST neurons and PV neurons receive closely matching numbers of incoming excitatory connections and have the same mean weight in our model, so the increase in SST responsiveness was likely not due to increased excitatory input (Fig. S8a). Instead, we found that SST neurons receive less inhibitory input from all other inhibitory neuron types in layer 2/3, such that SST neurons receive less inhibition than PV neurons in layer 2/3 (Fig. S8b-d). The overall lack of inhibitory input to SST neurons in layer 2/3 specifically was highlighted in a recent study on the reconstruction and analysis of the mouse connectome ^44^.

We then tested the contribution of intrinsic excitability of SST neurons by creating 30 different versions of the low hierarchical area model, each with a single SST neuron in layer 2/3 swapped with the single-neuron model for the most responsive PV neuron in layer 2/3. Our goal was to compare the response of SST neurons and PV neurons when embedded within the same circuit, and therefore receiving the same amount of excitatory and inhibitory input. We swapped a single SST cell for a single PV cell in each model, to mitigate potential network effects from changing many cells at once within a circuit. We then repeated the single-cell stimulation paradigm and analyzed the response of the PV neuron embedded in the circuitry of an SST neuron. We found that the swapped PV cells embedded into the SST cell’s location in the circuit were more responsive to the single-cell stimulation than the PV cells in the original model, confirming that the mean decrease in inhibitory input to SST cells contributed to the increased activity of SST cells in response to the single-cell stimulation paradigm (Fig. S8e). We also found that the swapped PV cells were less responsive than the SST cells in the original model, suggesting that the SST cell models are indeed more excitable than the PV cell models.

In summary, we found that the increased responsiveness of SST neurons was due to both intrinsic excitability and a relative lack of inhibitory input (Fig. S8). SST neurons may therefore contribute to maintaining stability in higher order cortical areas during periods of increased input to excitatory cells.

### Single-cell inhibitory stimulation is more depolarizing across the cortical hierarchy, regardless of stimulated cell type

We next repeated the single-cell stimulation paradigm, both experimentally and computationally, in inhibitory neurons in layer 2/3, instead of excitatory neurons. Inhibitory neurons were photostimulated in layer 2/3 of MOp while activity of the surrounding neurons was recorded with 2-photon imaging. The stimulation period lasted 25 ms. The effect of stimulating inhibitory neurons is broad local inhibition across all cells, up to approximately 200 µm from the stimulated neuron, both *in vivo* and in the low-hierarchy model, model 1 (Fig. 8b).

**Figure 8.**
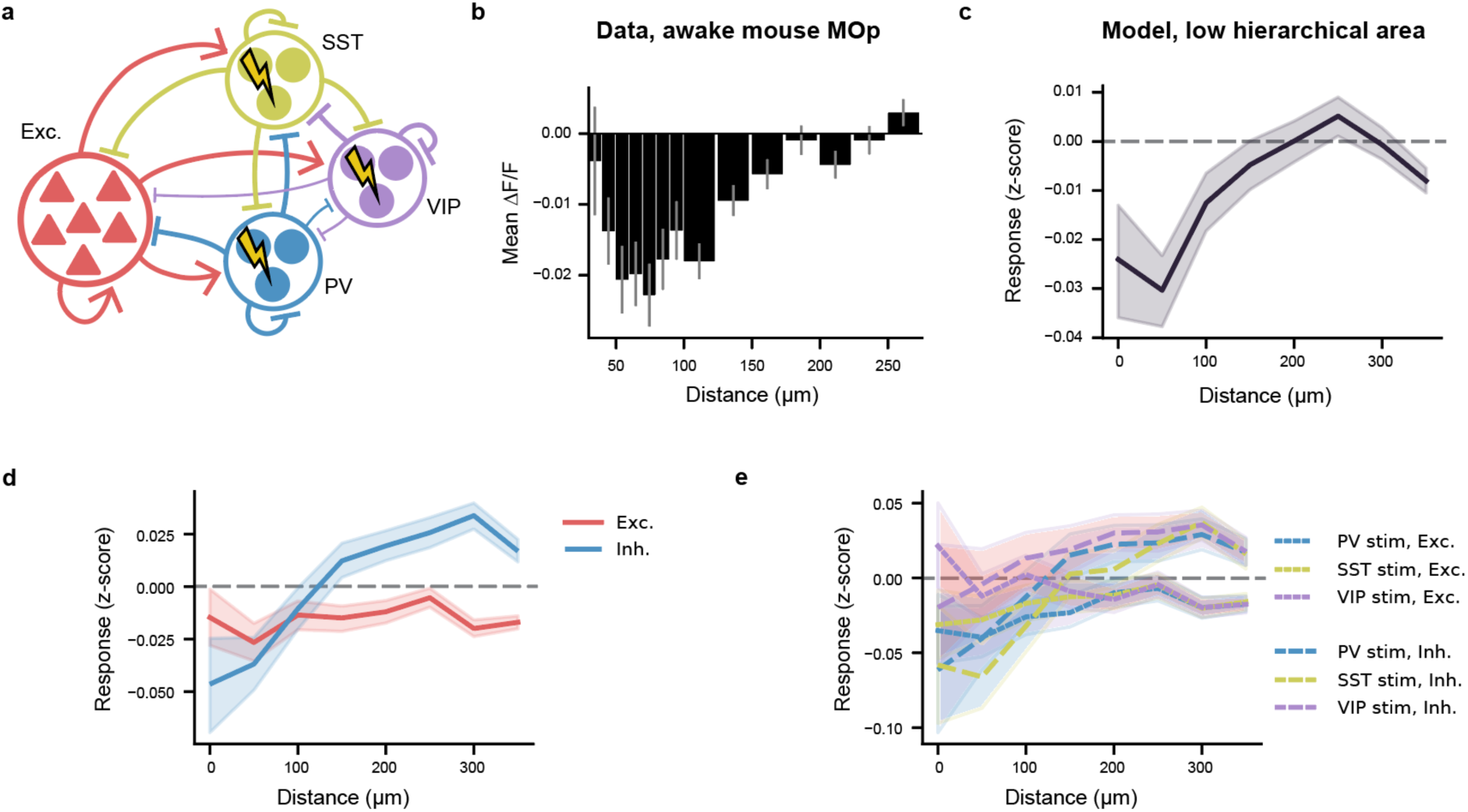
Single-cell stimulation of inhibitory neurons. a. Schematic of the single-cell inhibitory neuron stimulation experimental paradigm. b. *In vivo* response of neurons to single-cell stimulation of inhibitory neurons. c. *In silico* response of L2/3 model neurons to single-cell stimulation of inhibitory neurons d. *In silico* response of L2/3 model neurons to single-cell stimulation of inhibitory neurons, separated by excitatory and inhibitory neurons. e. *In silico* response of L2/3 model neurons, separated by excitatory neurons (solid lines) and inhibitory neurons (dashed lines) and stimulated neuron type.

In the models, we stimulated 2-4 PV, SST, and VIP neurons, one cell at a time, for 25 trials in each model (Fig. 8a). We observed local inhibition of the surrounding neuron up to approximately 200 µm, matching the experimental results (Fig. 8c).

We then separated the responses by cell type in the simulation results. On average, excitatory cells were broadly inhibited by the stimulation paradigm, while inhibitory cells were inhibited within 100 microns of the stimulated neuron, and disinhibited further than 100 microns away (fig. 8d). PV and SST neuron stimulation led to similar response profiles in both excitatory and inhibitory neurons in layer 2/3, while VIP stimulation led to weak disinhibition of excitatory cells within 50 microns of the stimulated VIP neurons (fig. 8e).

## Discussion

We created biologically-realistic models of circuits across the rodent cortical hierarchy and simulated responses to a range of standard experimental paradigms. These models systematically integrate observations about differences in local circuit composition and architecture across cortical areas, allowing us to examine how these structural features contribute to circuit function across the hierarchy. While many mechanistic and theoretical models of cortical circuits have focused on individual layers, particularly layer 2/3, our models incorporate all major cortical layers and their interactions within the local circuit. In addition, by including multiple diverse excitatory and inhibitory cell types, our models allow us to systematically examine how changes in the composition of these cell types influence circuit function across the cortical hierarchy. Other biological elements of the circuits, such as neuron and synapse type models or connectivity were derived systematically from experimental cellular and synaptic physiology measurements and connectomic data, providing biologically realistic constraints for our exploration of the circuit properties across the cortical hierarchy. Cells with similar tuning preferences were preferentially connected and also had higher weights ^57^; see ^45^ for more details.

Our simulations show that, while firing rates and activity patterns are qualitatively similar across the models of cortical areas, we find significant gradients in the underlying excitatory/inhibitory balance, leading to increased stability in response to perturbation as one moves from lower to higher levels of cortical hierarchy. Additionally, we find that increasing the density of SST neurons increases the prevalence of convergent motifs in higher order cortical models, suggesting that variation in SST abundance may contribute not only to the regulation of circuit stability but also to how information is integrated within local cortical networks. Finally, we also find that SST neurons are especially important for maintaining stability in response to perturbation in layer 2/3.

Lower cortical hierarchy areas are more sensory and associated with a more feedforward network structure and processing of mainly a single type of information, while higher cortical hierarchy areas are more associated with a convergent network structure and have a role in integrating multiple sources of information ^50^. To this end, higher order cortical areas must receive many inputs while maintaining E/I balance and the ability to perform complex cognitive tasks, such as working memory and decision-making. Here we provide evidence that increasing SST neuron density may help integrate information by increasing prevalence of convergent motifs in a circuit and maintaining balance in response to increasing external input. Our results demonstrated that single-cell stimulation leads to a network-wide shift in excitatory/inhibitory balance, and SST neurons were necessary to provide additional inhibitory input in higher order cortical regions.

The responsiveness of SST neurons to excitatory stimulation was due to both increased excitability of SST cells and a relative lack of inhibitory input to layer 2/3 SST neurons. This lack of inhibitory input to layer 2/3 SST neurons is a main finding of the MICrONS Connectivity dataset, which demonstrates that VIP neurons, the main source of inhibition to SST neurons, have sparse output to SST neurons in layer 2/3 ^44^. Of note, short-term plasticity between excitatory and SST neurons is potentiating and especially strong in layer 2/3 ^34^, providing a third mechanism for the strong response of SST cells to external stimulation that was out of scope of our model. Overall, our findings replicate previous single-cell stimulation experimental results ^73,76^ and may relate to findings that layer 2/3 SST neurons play a role in cortical stability and synchrony ^77,78^.

We found that single-neuron timescales slightly decreased across the cortical hierarchy rather than increased, suggesting connectivity and SST neuron density are not the mechanism behind the experimentally observed increasing timescales across the hierarchy. Other potential mechanisms are long-range connectivity ^55^, neuromodulator input ^21,22^, increasing receptive field size ^33^, or variations in subcortical input. The working model is that longer timescales are a global, rather than local, property of the brain, supported by the finding that timescale distributions follow the same power law distribution across all brain regions and vary with task ^54,56^.

It is important to note that here we modeled the cortical hierarchy as a one-dimensional gradient from sensory to higher order cortical areas, but multiple other cortical organization schemes have been proposed. There is, for example, structure as well along dorsal and ventral pathways of the cortex, and the motor cortex, while considered to be low in hierarchy like primary sensory regions, is also the main output region of the brain. More generally, the present models focus on local cortical circuits and therefore do not capture the full contribution of long-range corticocortical, thalamocortical, and other subcortical inputs to the differences in circuit dynamics across cortical areas.

Incorporating these long-range connections will be important for determining how local circuit properties interact with inputs from other brain regions and for understanding whether some of the observed functional gradients emerge only at the level of larger distributed networks. Future studies could also extend the models to include a broader diversity of cortical areas and more detailed variation in neuronal and synaptic properties, as additional experimental data become available. Similarly, examining a wider range of stimuli, behavioral states, and tasks will provide opportunities to test whether the relationships between local circuit structure, stability, and information integration identified here generalize across different modes of cortical function. Thus, the present models provide a framework that can be progressively expanded as structural, physiological, and functional data become available across the brain and as simulation capabilities develop.

In conclusion, our modelling results suggest an overall role of local circuits in integrating external information and maintaining stability across the cortical hierarchy. The systematic incorporation of differences in cortical cell-type composition, connectivity, and architecture allowed us to identify relationships between these structural features and circuit function, including a particularly important role for SST neurons in maintaining stability and shaping convergent network motifs in higher-order cortical areas. At the same time, our results indicate that some functional properties, such as the increase in neural timescales across the cortical hierarchy, likely depend on factors beyond the local circuit. Together, these findings illustrate how biologically realistic large-scale modeling can provide a framework for connecting increasingly detailed structural and physiological measurements to the functional properties of cortical circuits. To facilitate further investigation and enable the community to build on this work, all models, simulation results, and code developed in this study will be made publicly available upon publication.

## Methods

The models used in this study were based on the models of primary visual cortex created in ^39,45^. The models represent a column with a 400 µm radius in primary visual cortex, with 5 layers (layer 1, 2/3, 4, 5, and 6) and 200 different cell models representing 19 cell types (PV, SST, VIP, and excitatory cell types in each layer), based on the Allen Mouse Brain Atlas ^79^ (https://atlas.brain-map.org). The individual cell models were fit to *in vitro* measurements^46^ and are publicly available via the Allen Cell Types Database (https://celltypes.brain-map.org/). The connections between each cell type are based on both the mapped connectome^38^ and a multi-patch synaptic physiology dataset^34^. The weights are based on the same multi-patch synaptic physiology dataset, then adjusted to match the firing rates from neuropixels recordings in the visual cortex with opto-tagged excitatory, PV, SST, and VIP neurons ^33^. Input to the cortical model is from a filter-based model^80^ of the lateral geniculate nucleus (LGN) of the thalamus, with units fit to 14 cell types in the thalamus to represent experimentally-observed diversity in spatiotemporal filtering characteristics ^81^. The LGN units represent visual input on a 2D plane in visual space, 240 degrees (horizontal) by 120 degrees (vertical). Connections from the LGN filter units to VISp were distributed retinotopically to preserve the spatiotemporal structure of input to the model. The model of the visual cortex is available on https://brain-map.org/our-research/computational-modelling and more details are available in ^39,45^.

We created 12 additional models by varying either the connection density, the inhibitory neuron type ratio, or both. Each set of new models was based on data for primary motor cortex, parietal cortex, anterior cingulate cortex, or ventral orbitofrontal cortex. The connection probability in each model is described as a gaussian distribution, with maximum probability of connection, *p_max_* and standard deviation *σ*. To increase the connection density, we increased *p_max_* proportional to the decrease in cell density in each model, and then varied *σ* until the number of synapses in the model was consistent with the number of synapses in the base model. This was done to facilitate adjusting the weights in each model. The weights were adjusted so that the mean total of the number of connections multiplied by each weight was consistent for each neuron population. We constructed changes in the inhibitory neuron density by adjusting the ratio of PV, SST, and VIP neurons, following the data in ^25^, which includes data from ^24^.

The standard stimulus presented to each model was a drifting grating input (2.5 seconds duration, 2 Hz temporal frequency, 80% contrast, 0.04 cpd, at 8 different angles from 0 to 315 degrees). A drifting grating was chosen as a stimulus for several reasons: its ubiquity as an experimental stimulus^33^, validation in previous versions of the model^39,45^, and to facilitate direct comparison between models. Spontaneous firing rates were calculated as the number of spikes during the first 500 ms of a trial, divided by 0.5 s. The maximum mean rate was calculated as the maximum firing rate during stimulus presentation.

Stimulus selectivity was measured as the ratio of difference of response at the preferred stimulus (*R_pref_*) to the response at the least-preferred stimulus (*R_ort_*_ℎ*o*_), to the sum of the responses to the preferred and least preferred stimulus.

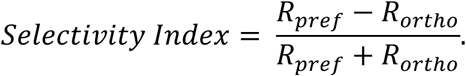

Input to each cell, *i*, was measured as the sum of the number of spikes per second (*N_spikes_*_,*j*_) of each input cell, *j*, multiplied by the weight of the synapse from that cell, *W_ij_*:

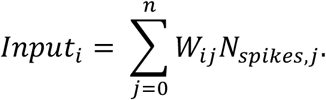

The value ß was calculated as the ratio of all excitatory input to the total input ^48^:

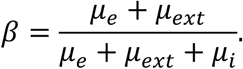

We calculated the time constant by first binning the spikes of all excitatory neurons with a mean firing rate greater than 0.005 Hz were binned into 50 ms bins. We then fit an autoregressive (AR) model to the binned spikes of each neuron^53,54^. The time constant, *τ*, was calculated from the eigenvalues, *λ*, of three AR coefficients (*a*^1^, *a*^2^, *a*^3^) as follows:

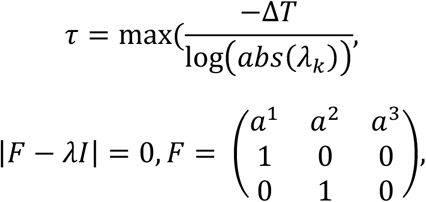

where Δ*T* is the time bin size (50 ms).

To calculate the graph properties of each network, we converted the middle 200 µm of each layer in each model to a networkx graph in python. We used the middle 200 µm for computational feasibility. The clustering coefficient, *c*, was calculated using the algorithm for an unweighed graph in networkx, as:

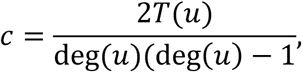

where *T*(*u*) is the number of triangles through node *u* and deg(*u*) is the degree of *u*.

The number of motifs in each network was calculated by counting the number of each motif within the middle 200 microns of each layer in each network. We converted each network into a networkx graph and used the python package dotMotif^82^ to count the number of each motif. We then generated 200 Erdos-Renyi random null graphs for each model, matched for the number of nodes and edges, and measured the number of each motif in each graph to get a null distribution. The over-representation score for each motif was calculated as the z-score of the number of motifs in the connectome-based models versus the node-and edge-matched null distribution.

We determined whether a network was an inhibition-stabilized network by providing a simulated current clamp input to all inhibitory neurons in the network at increasing intensity and measuring the change in firing rate of inhibitory and excitatory neurons. We then measured whether the excitatory input to inhibitory neurons decreased because of increased inhibitory stimulation, to ensure that the mechanism of decreased inhibitory activity was a decrease in excitatory input.

The single cell stimulation paradigm was simulated in each model by providing a simulated depolarizing current clamp input to one neuron at a time in layer 2/3 of each model, for 25 trials. We selected neurons within 30 microns of the center of the model, to ensure we could measure the response of neurons up to 200 microns away. There was a 1 second rest period in between stimulation. To take into account the baseline activity of each neuron, the normalized response of each neuron, *R_norm_*, was measured as:

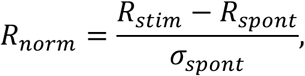

where *R_stim_* is the activity of the neuron during the stimulation period, *R_spont_* is the activity of the neuron during a control trial with no stimulation, and *σ_spont_* is the standard deviation of the neuron’s activity during the spontaneous period.

The additional input to each excitatory neuron as a result of stimulation was measured in the same manner as above.

The excitatory/inhibitory balance was measured as the ratio of mean of the excitatory firing rates in the model divided by the mean of the inhibitory firing rates in the model.

