## Supplemental figure 1 for "Gradients of local circuit architecture shape stability and information integration across the cortical hierarchy"

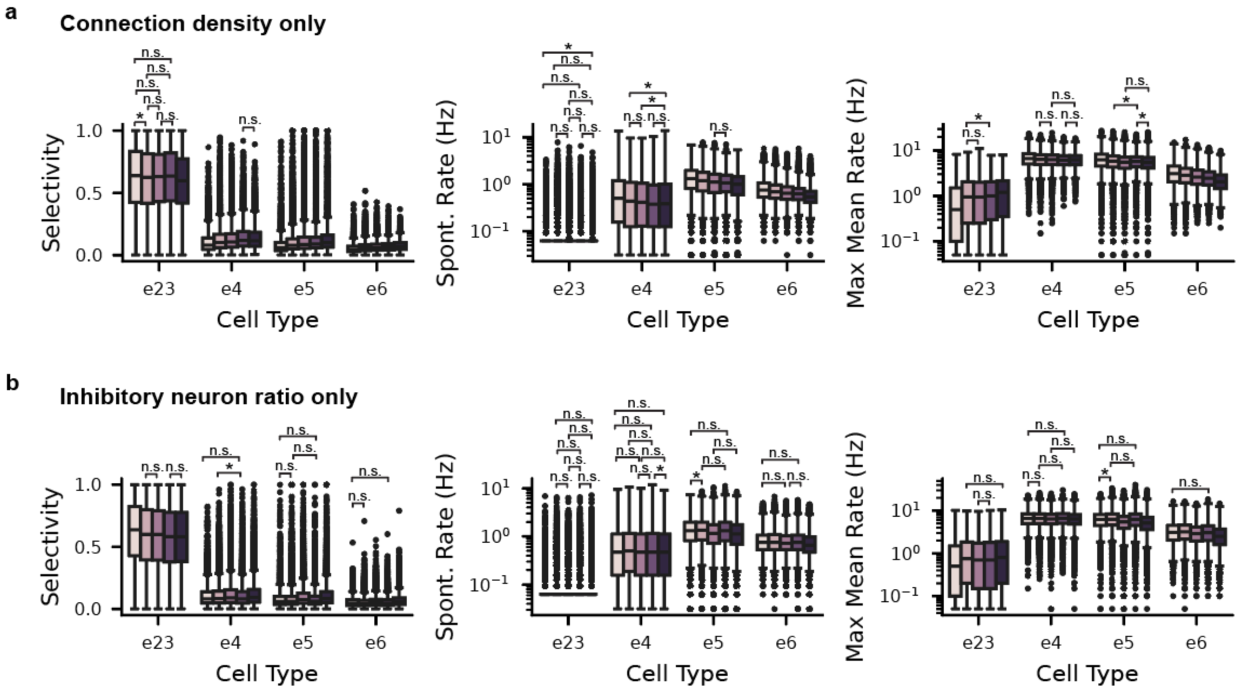


Supplemental figure 1: Population characteristics of models with changes in connection density or inhibitory neuron ratio.

a: Stimulus selectivity, (left), spontaneous firing rate (middle), and maximum mean firing rate during stimulus presentation (right) of excitatory cells across layers and across models with increasing connection density.

b: same as (a), but for models with changing inhibitory neuron type ratio.
