## Supplemental figure 2 for "Gradients of local circuit architecture shape stability and information integration across the cortical hierarchy"

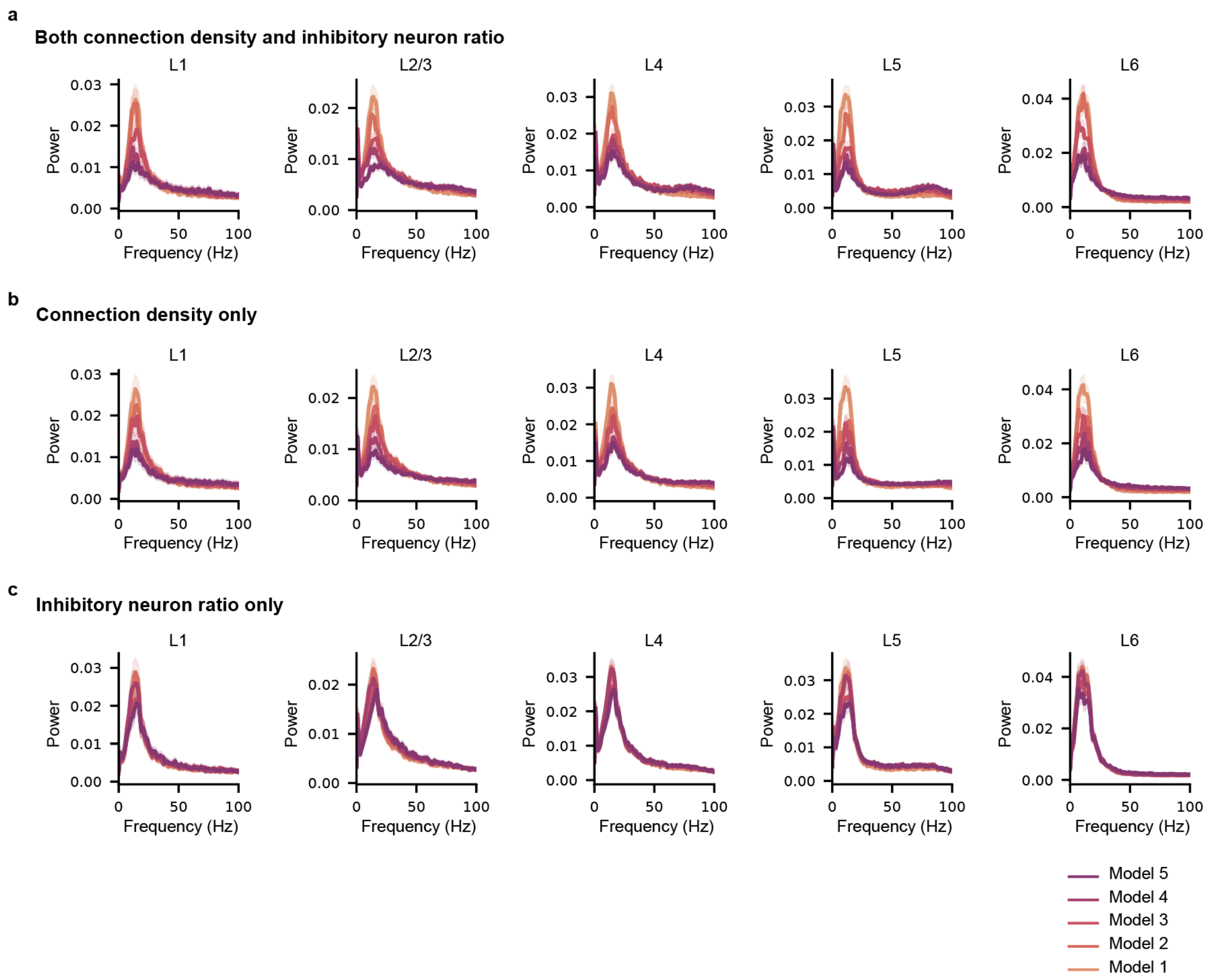


Supplemental figure 2: Power spectrum densities across layers and cortical regions.

a: Power spectrum densities of mean population firing rates in each layer in models across the cortical hierarchy, normalized by the total sum of the power spectrum density.

b, c: Same as (a), but for models with only changes to connection density (b) or inhibitory neuron subclass density (c).
