## Supplemental figure 3 for "Gradients of local circuit architecture shape stability and information integration across the cortical hierarchy"

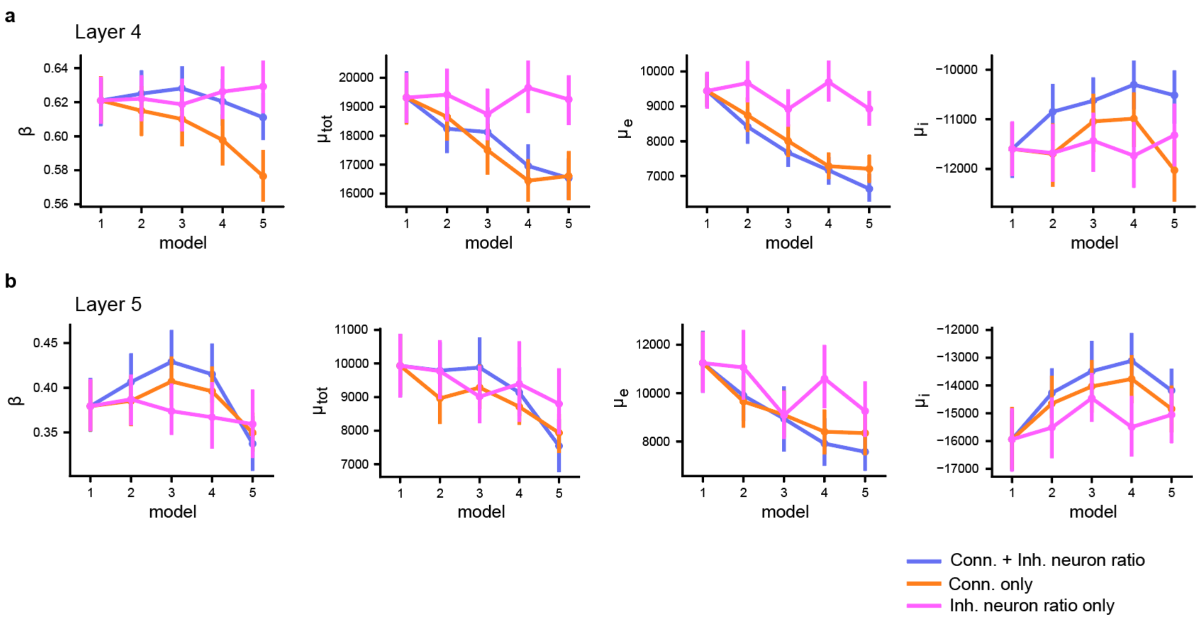


Supplemental Figure 3: Input to excitatory cells in layers 4 and 5 across the cortical hierarchy.

a: input metrics of excitatory cells in layer 4. From left to right: ß, total input ($\mu_{tot}$), total excitatory input ($\mu_e$), and total inhibitory input ($\mu_i$).

b: Same as (a) but for layer 5.
