## Supplemental figure 4 for "Gradients of local circuit architecture shape stability and information integration across the cortical hierarchy"

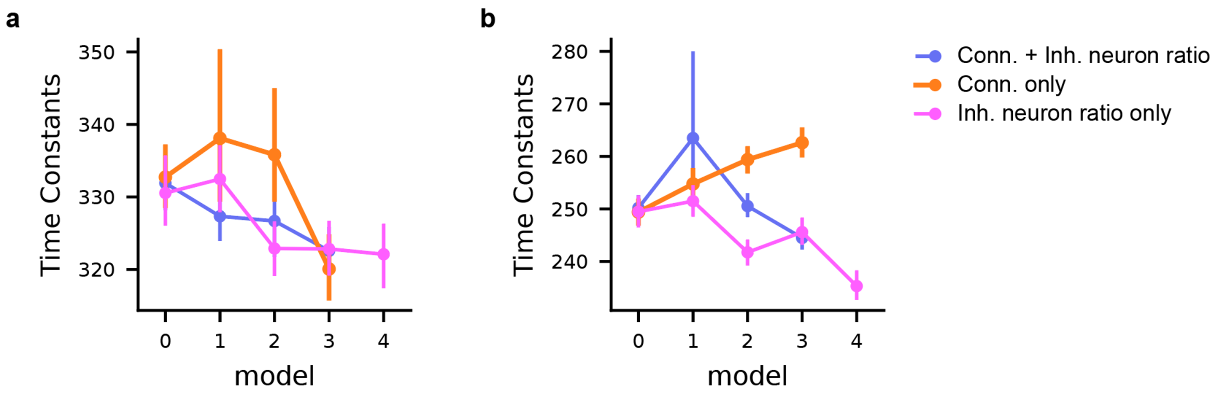


Supplemental 4. Time constants across the cortical hierarchy in layers 4 and 5.

a. Time constants in layer 4 in all models across the cortical hierarchy.

b. Same as (a) but for layer 5.
