## Supplemental figure 5 for "Gradients of local circuit architecture shape stability and information integration across the cortical hierarchy"

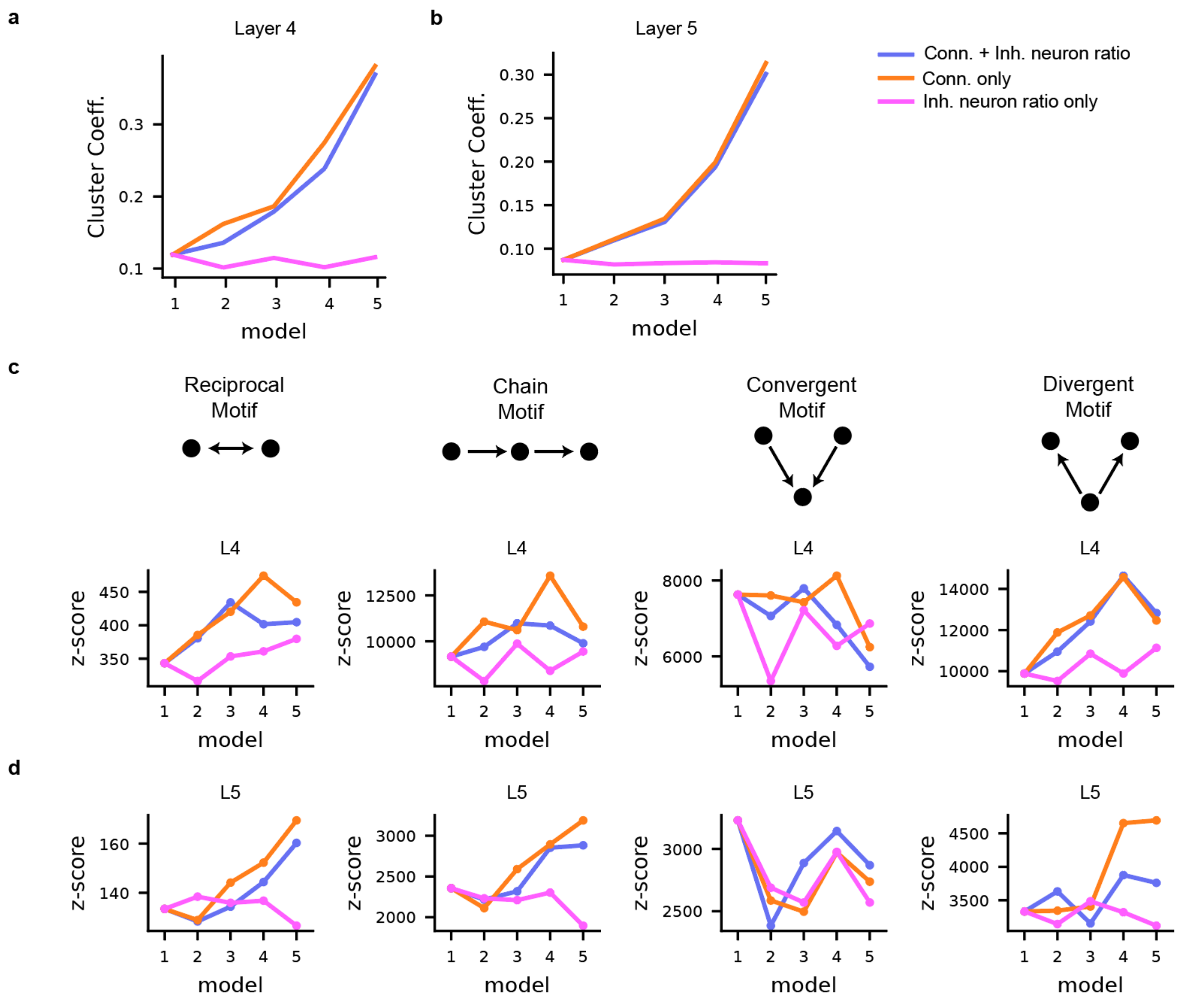


Supplemental figure 5. Graph metrics of layers 4 and 5

a. Cluster coefficient in layer 4 of all models across the cortical hierarchy.

b. Same as (a) but for layer 5.

c. Motif overexpression in layer 4 of all models across the cortical hierarchy. Top: representation of each motif type. Bottom: motif overexpression score.

d. Same as (c) but for layer 5.
