## Supplemental figure 6 for "Gradients of local circuit architecture shape stability and information integration across the cortical hierarchy"

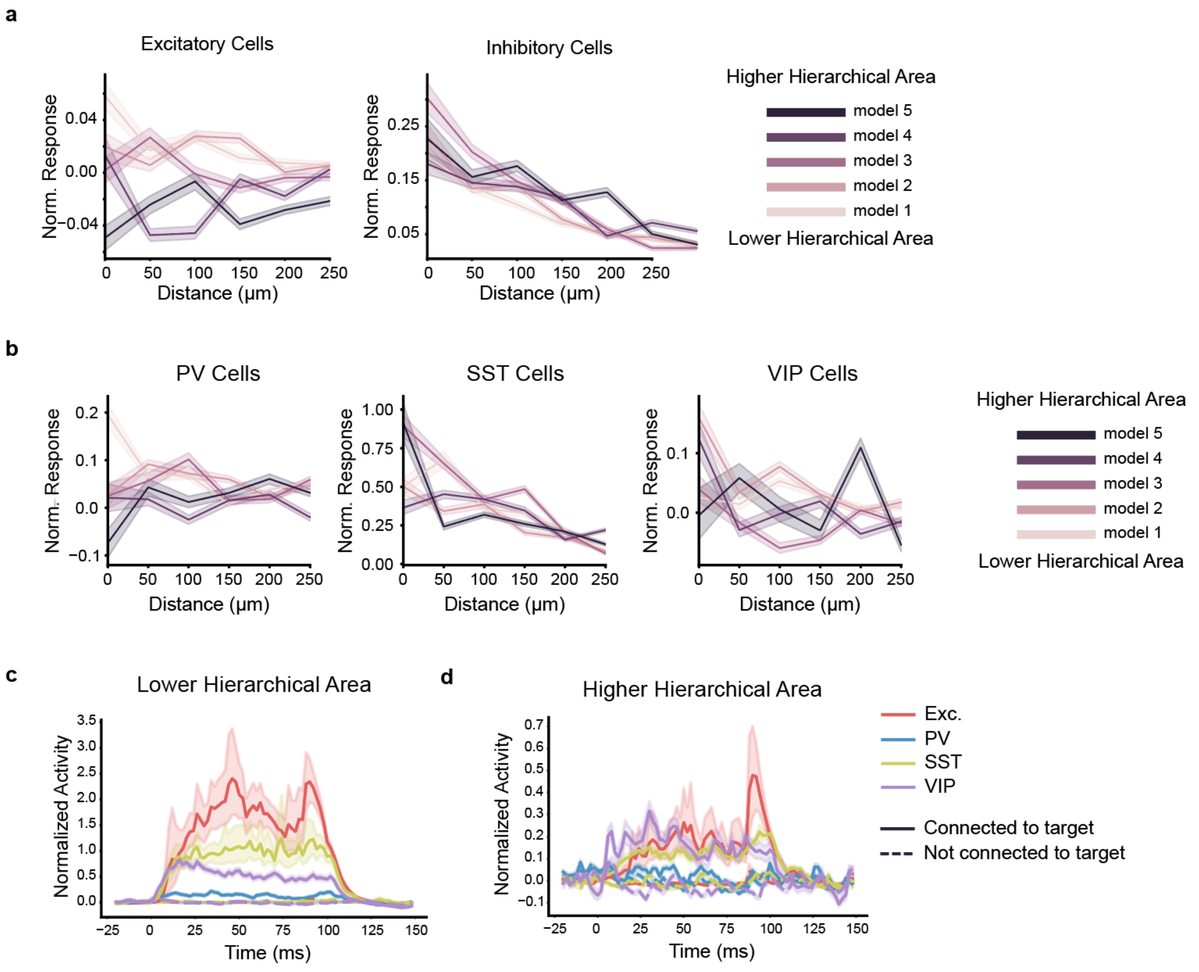


Supplemental figure 6: Response to single-cell stimulation across the cortical hierarchy.

a: Normalized response of excitatory (left) and inhibitory (right) cells at increasing distance to the stimulated neuron. Responses are normalized to each cell's firing rate during spontaneous activity. Darker colors denote higher order cortical models.

b: Same as (a) but grouped by inhibitory neuron subtypes: PV (left), SST (middle), and VIP (right).

c, d: Time course of responses to the single-cell stimulation paradigm, grouped by neuron subtype (Exc., PV, SST, or VIP) and whether a neuron is directly downstream of the stimulated neuron (solid lines) or not connected to the stimulated neuron (dashed lines), for a lower order cortical model (left) and a higher order cortical model (right).
