## Supplemental figure 7 for "Gradients of local circuit architecture shape stability and information integration across the cortical hierarchy"

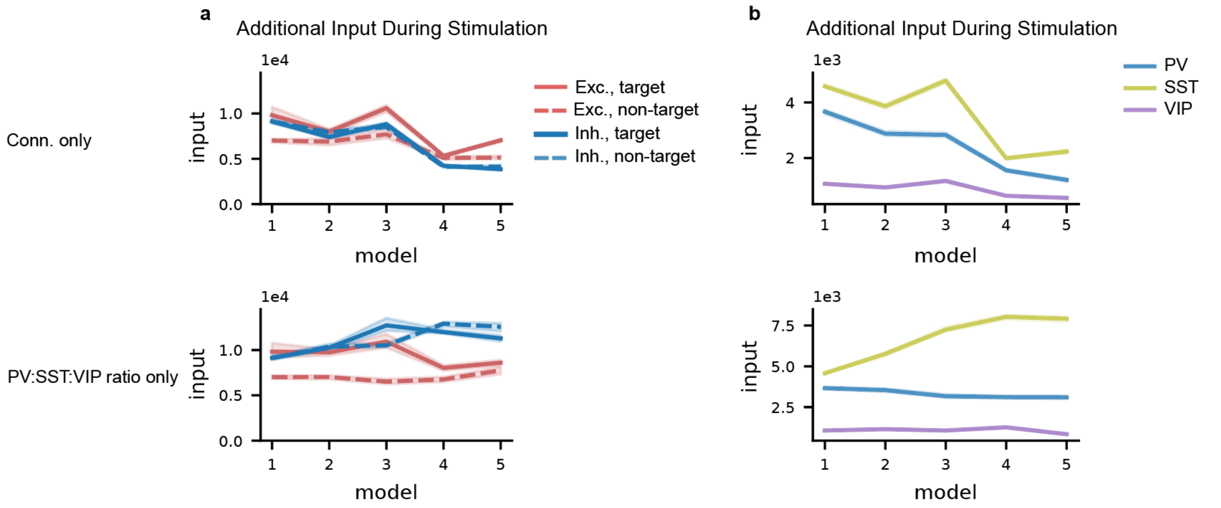


Supplemental figure 7: Input to excitatory cells during the stimulation paradigm.

a: Excitatory (red) and inhibitory (blue) input to layer 2/3 excitatory neurons, grouped by target (solid) and non-target (dashed) neurons. Top: models with increasing connection density; bottom: models with increasing SST neuron density.

b: Inhibitory subtype input to layer 2/3 excitatory neurons. Blue: PV input; yellow: SST input; purple: VIP input. Top: models with increasing connection density; bottom: models with increasing SST neuron density.
