## Supplemental figure 8 for "Gradients of local circuit architecture shape stability and information integration across the cortical hierarchy"

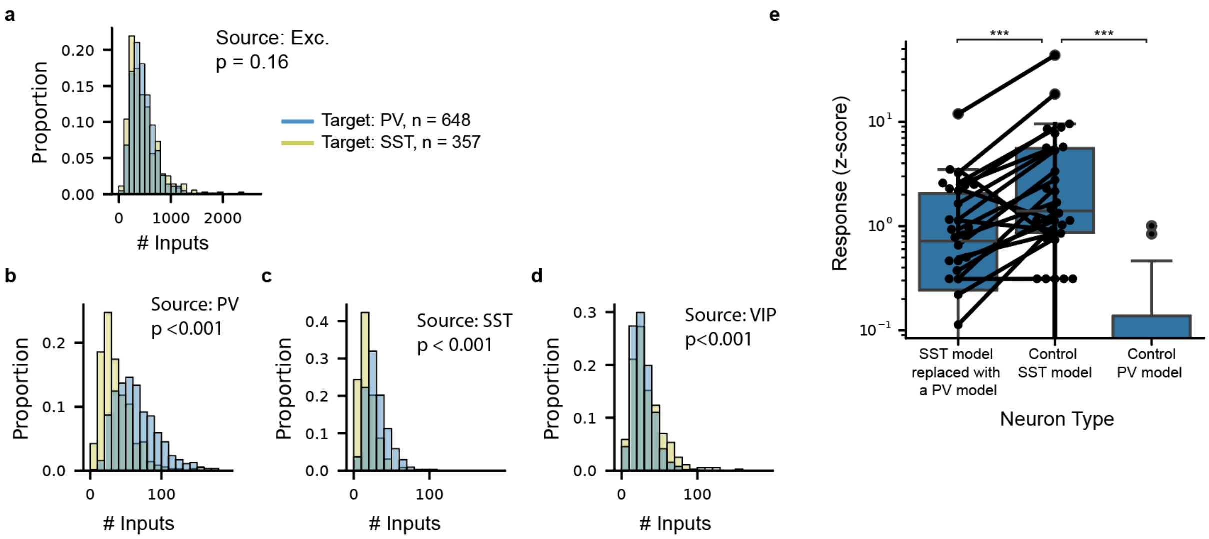


Supplemental figure 8: Layer 2/3 SST neuron responsiveness is due to low inhibitory input and high intrinsic excitability, rather than high excitatory input.

a: Distribution of the number of excitatory inputs to PV (blue) and SST (yellow) neurons in the model.

b-d: Distribution of the number of PV inputs (c), SST inputs (d), and VIP inputs (e) to PV and SST neurons in the model.

e: Response of a PV neuron model swapped with an SST neuron, such that the PV neuron is embedded into the circuitry of an SST cell, repeated for 30 different SST cells (left), response of SST neurons within the circuit (middle), and the most responsive PV neuron model (right) during the single-cell stimulation paradigm.
