## Supplemental figure 9 for "Gradients of local circuit architecture shape stability and information integration across the cortical hierarchy"

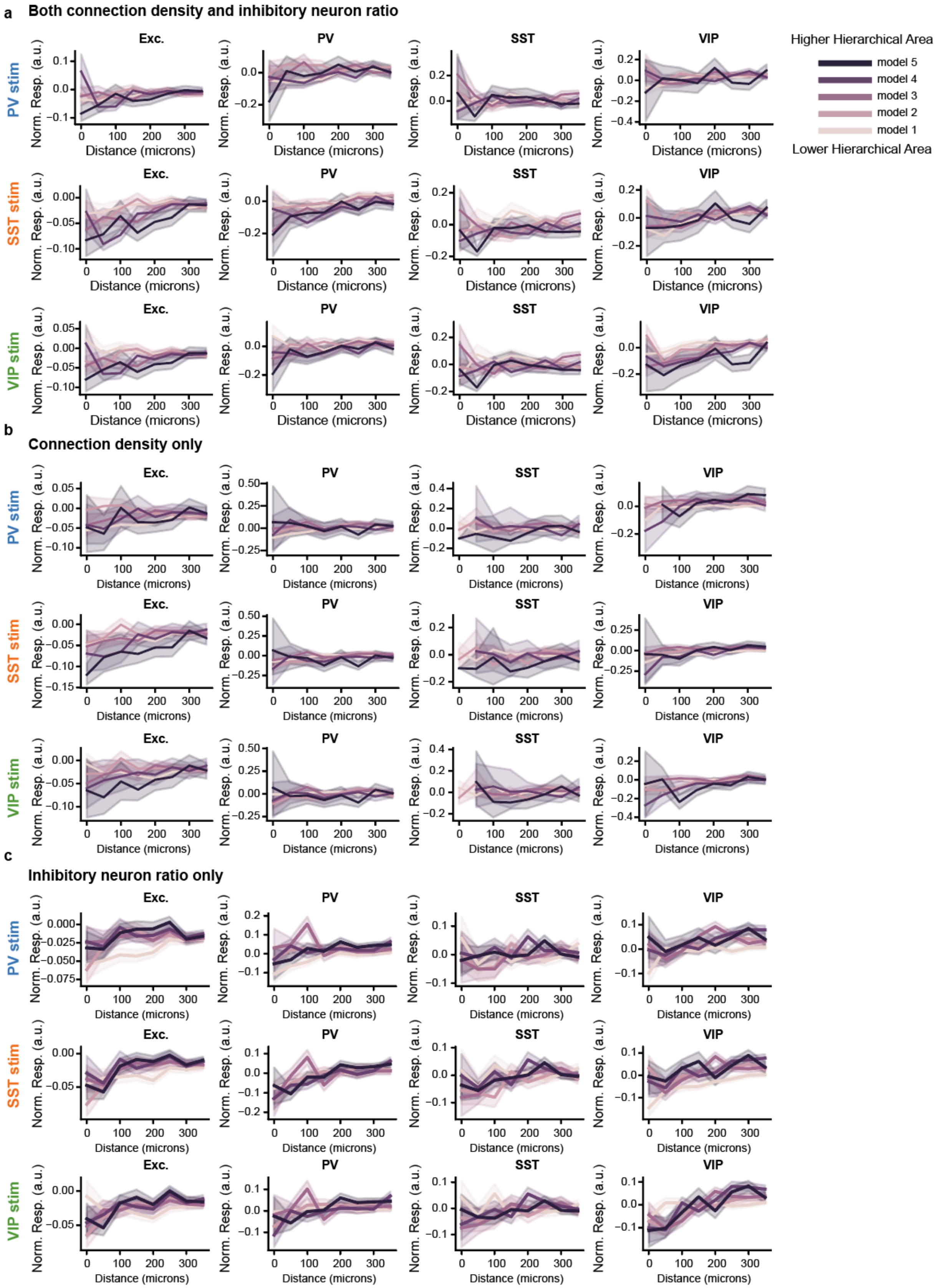
a
Both connection density and inhibitory neuron ratio
b
Connection density only
c
Inhibitory neuron ratio only
PV stim
SST stim
VIP stim
Exc.
PV
SST
VIP
Norm. Resp. (a.u.)
Distance (microns)
Higher Hierarchical Area
model 5
model 4
model 3
model 2
model 1
Lower Hierarchical Area

Supplemental figure 9: Response to single-cell inhibitory stimulation.

a: Normalized response of (from left to right) excitatory, PV, SST, and VIP cells at increasing distance to a stimulated inhibitory neuron. Responses are normalized to each cell's firing rate during spontaneous activity. Darker colors denote higher order cortical models. Top: responses to PV neuron stimulation; middle: responses to SST neuron stimulation; bottom: responses to VIP neuron stimulation.

b, c: Same as (a), but for the models with changes in connection density only (b) and inhibitory neuron ratio only (c). Due to the lower fraction of inhibitory neurons in the models, in some higher order models with reduced neuron density, there is no data in the bin closest to the stimulated neuron.
